# Comparative co-occurrence network analyses of the cichlid fish gut microbiota: community conservation and diet-associated shifts

**DOI:** 10.1101/2020.05.26.118232

**Authors:** Joan Lluís Riera, Laura Baldo

## Abstract

Co-occurrence networks of bacteria associations are a powerful approach to explore ecologically relevant aspects of the gut microbiota structure, beyond community composition alone. Here we exploit the remarkable diversity of cichlid fishes and their multiple lake assemblages to investigate a) network features and patterns of microbial associations that were robust to major phylogeographical variables, and b) community structure changes along cichlid dietary shifts. We tackled these objectives using the large gut microbiota sequencing dataset available (nine lakes from Africa and America), building geographical and diet-specific networks and performing comparative analyses. Major findings indicated that lake and continental networks were highly resembling in global topology and node taxonomic composition, suggesting important constraints in the cichlid gut community assembling. A small fraction of the observed co-occurrence pairwises was conserved across all lake assemblages; while the origin and ecological relevance of these core associations remains unclear, their persistence suggests a potential functional role in the cichlid gut. Comparison of carnivores and herbivores-specific networks as well as mapping of diet-specific values on the African Lake Tanganyika network revealed a clear community shift as a function of diet, with an increase in complexity and node taxonomic diversity from carnivores-omnivores-plantkivores to herbivores. More importantly, diet-associated nodes in herbivores formed complex modules of positive interactions. By intersecting results from association patterns and experimental trials, future studies will be directed to test the strength of these microbial associations and predict the outcome of community alterations driven by diet.

**Importance:** The gut microbiota is a complex community of interacting bacteria. Predicting patterns of co-occurrence among microbes can help understanding key ecological aspects driving community structure, maintenance and dynamics. Here we showed a powerful application of co-occurrence networks to explore gut bacteria interactions in a primary model system to study animal diversification, the cichlid fishes. Taking advantage of the large scale of phylogeographical and ecological diversity of this fish family, we built gut microbiota networks from distinct lake and continental fish assemblages and performed extensive comparative analyses to retrieve conserved and trait-specific patterns of bacteria associations. Our results identified network features that were independent from the fish biogeography and that indicated an important host selection effect on gut community assembling. Focusing on a single lake assemblage, and therefore excluding the major geographical effect, we observed that the gut microbiota structure dramatically shifted from carnivore to herbivore fishes, with a substantial increase in the number and complexity of microbial interactions.

## Introduction

Compositional aspects of the gut microbiota (e.g. number of distinct Operational Taxonomic Units – OTUs - their relative abundance and phylogenetic relationships) reflect that fraction of the environmentally available microbes that are able to colonize the gut and assembled into a community. It is known the host intestinal environment can actively shape these microbial communities by imposing strong constraints in the colonization and by mediating community assembling through specific niche offering (Rawls et al., 2006)(Yan et al., 2016)(Burns et al., 2016). The combined influence of the host and the environmental mediated factors (e.g. dietary inputs) can therefore result into specific and to some extent predictable gut communities (Baldo et al., 2017)(Ley et al., 2008)(Foster et al., 2017). Still, the extent to which deterministic rather than stochastic processes guide microbes co-existence and ultimately their assembling into a community remains a matter of debate (Sieber et al., 2019)(Burns et al., 2016). Unclear is also whether patterns of microbial co-occurrence are driven by specific strain/OTUs or more generally guided by conserved features at higher bacterial taxonomic levels (from genus to phylum-levels), therefore assuming functional equivalence among similar taxa.

A powerful approach to start exploring the forces that control gut microbial community structure and its stability/dynamics is through co-occurrence networks (Röttjers & Faust, 2018). This increasingly used analytical tool relies on microbial abundance data obtained from extensive sequencing data (typically a matrix of OTUs) to infer microbe interactions as a function of their covariation patterns across a wide number of samples (Faust & Raes, 2012). Association patterns are deduced through a number of different correlation metrics, while significance is typically assessed through permutations, with the assumption that a non-random pattern is shaped by niche processes driving coexistence; these include ecological-based interactions, with positive associations putatively indicating cross-feeding or partial niche overlap, and negative associations indicating competitive exclusion or predation (Faust & Raes, 2012) (Williams et al., 2014).

Comparative analyses of microbial networks built from distinct datasets that vary at one or multiple sample traits (either host or environmental) are particularly powerful to explore microbial community dynamics (Faust et al., 2015). This type of approach can retrieve network commonalities across systems, e.g. microbial associations that are persistent along major host/environmental-associated variables (i.e. taxonomical, spatial, temporal and ecological), as well as unveil system and trait-specific co-occurrence patterns (e.g. with the host health status, ethnicity and dietary habits) (Jackson et al., 2018)(Mandakovic et al., 2018)(Faust et al., 2015)(Hegde et al., 2018)(Li et al., 2019). While not being conclusive in terms of inferences, these methods allow a first exploration beyond microbiota compositional aspects and set the ground for empirical testing of novel hypotheses (Röttjers & Faust, 2018).

Recent studies have successfully applied this comparative approach to the exploration of microbial associations at distinct scales of biological organization, i.e. across host populations (Jackson et al., 2018), species (Hegde et al., 2018) and within and across distinct biomes (Mandakovic et al., 2018)(Williams et al., 2014)(Faust et al., 2015). Their application to studies of animal-associated microbiotas remains, however, relatively scarce and has primarily targeted humans, along with few other vertebrates (e.g., see (Jackson et al., 2018)(Li et al., 2019)). Particularly, little is known about how much of the observed gut microbial diversity engage into robust, conserved interactions in nature, and whether these interaction patterns are maintained or modified during the natural process of host adaptation, following both phylogeographical and ecological diversification. A main question is whether and how the microbiota community network adjusts to dietary shifts that might occur among closely related host species.

Cichlid fishes provide an attractive system to investigate gut microbe-microbe association patterns and community changes during host divergence. Cichlids represent an iconic fish family (Cichlidae), widely distributed across lakes and rivers in the subtropical/tropical regions and have served as a primary model to study speciation and rapid phenotypic diversification (Salzburger, 2009, 2018). The extraordinary range of ecological niches they occupy, even within highly reduced water pools (e.g. small crater lakes), are primarily driven by resource partitioning and niche displacement, following competition for local resources (Seehausen, 2006)(Muschick et al., 2012)(Barluenga et al., 2006)(Kautt et al., 2018). In the African Great Lakes (Tanganyika, Malawi and Victoria), repeated explosive adaptive radiations have led to the greatest levels of cichlid specialization in terms of morphology, behavior and dietary preferences (Curry-Lindahl et al., 1976)(Salzburger, 2018). Central American lakes, on the other hand, host more recent and less ecologically diverse fish assemblages (Kautt et al., 2018). While very diverse and widely distributed, cichlids are nonetheless a relatively young family (most species have evolved within the past 0.5 Mya,), with distinctive and unifying phenotypic traits (Salzburger, 2018), shaped by a common genomic structure (Brawand et al., 2015).

The cichlid ability to exploit a vast range of dietary inputs, along with several examples of trophic convergence across continental and lake assemblages, make this fish family a great model to study gut microbial co-occurrence patterns (Baldo et al., 2015, 2017, 2019)(Faber-Hammond et al., 2019). Recent analyses have shown a strong correlation between compositional aspects of the cichlid gut microbiota and dietary habits, suggesting a microbial role in the fish exploitation and optimization of feeding inputs (Baldo et al., 2017, 2019). These studies also revealed the existence of a small microbial component conserved across all cichlids (core). It remains unclear, however, whether these diet-associated and conserved compositional traits of the cichlid gut microbiota are driven by specific microbe-microbe interactions and how host ecology and geography influence these associations. Here we provided the first exploration of the cichlid gut microbial interactions through co-occurrence network analyses. We took advantage of a recently published large scale gut microbiota dataset (Baldo et al., 2019), including samples from two African and seven American lakes encompassing major dietary niches, to address the following major objectives: a) provide the first description of the cichlid gut community network; b) identify network features and pairwise occurrences robust to major host ecological and geographical variables (i.e. the core network); b) detect diet-specific association patterns that can help us understanding microbial community changes during cichlid dietary adaptation.

To reach these goals, we partitioned the dataset according to major geographic and dietary variables and built individual co-occurrence networks using a consensus of correlation measures. We then contrasted the networks generated to retrieve commonalities/differences in microbial associations across fish assemblages. Finally, we developed novel approaches to map the diet contribution to the network structure. Our results allowed to formulate interesting hypotheses on the constraints acting on the cichlid gut communities and their dynamics, setting the basis for future microbiome modeling and validation through empirical trials in this fascinating system.

## Results

### Comparative analyses of lake and continental networks

Here we used an ensemble approach of four correlation measures to infer robust gut bacterial association patterns in individual continental and lake fish assemblages. Specifically, we built co-occurrence microbial networks for the African and American datasets and for six individual lake datasets, including the two African lakes (Lake Tanganyika and crater lake Barombi Mbo) and four of the six American lakes (crater lakes Apoyo, Apoyeque and Xiloá and the large lake Nicaragua) (Fig. 1A-B). For the American lakes Masaya, Managua and Asososca León, sample numbers did not reach the minimum required (*n* = 20) for network building and sample contribution from these lakes was simply integrated into the American continental network.

**Figure 1:**
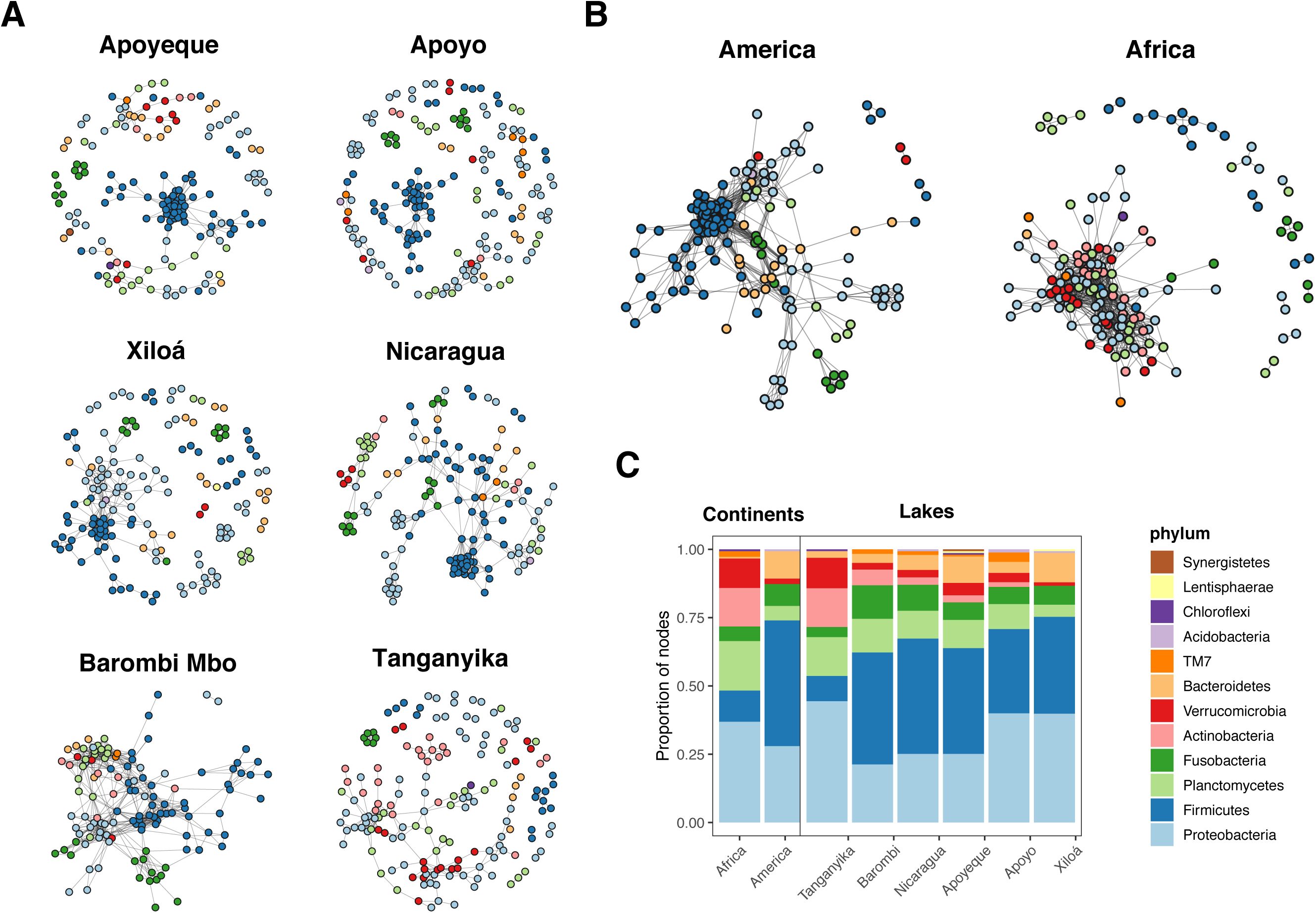
Co-occurrence network graphs of the cichlid gut microbiota and their node taxonomic composition. (A-B) Networks for individual lake assemblages (A) and continental datasets (B), where nodes are OTUs color-colored to the phylum level. (C) Proportion of nodes (OTUs) per phylum found in each geographical dataset (for the overall microbial taxonomic composition, see (Baldo et al., 2019), while for proportion of OTUs per phylum in the input dataset see Fig. S3). All American networks largely resembled in topology and node taxonomic content. Note the dense cluster of Firmicutes nodes present in all lake networks, but L. Tanganyika, encompassing the two families Clostridiaceae and Peptostreptococcaceae (see Fig. S1 for family level). Network interactions were inferred based on concordance among four co-occurrence measures in the ensemble package CoNet (Li et al., 2019) and visualized with *igraph* (Csardi & Nepusz, 2006).

All geographical-based networks were significantly distinct from a random graph of equal size (same number of nodes and edges), as calculated under the Erdos-Renyi model, and appeared to follow a scale-free distribution, where the node degree distribution fits a power-law (y = ax^b^), with correlation values for Africa = 0.877 (R^2^ = 0.491), and for America = 0.634, (R^2^ = 0.456) (Fig. 1A-B). Despite the heterogeneity of the sample data in terms of species phylogeny and ecology (42 species and five major diets, for details see (Baldo et al., 2019)), both individual lake and continental networks were largely comparable in size (number of nodes varying between 122 and 175) and global topological features (Table 1). OTUs engaging into significant associations (p < 0.05, BH correction) represented between 21% and 43% of the total OTUs per dataset present in the original input matrix (Table 1). The large majority of the associations (edges) were positive (copresence) for all nets, which were relatively poorly dense (0.02-0.10, graph density) and showed comparable clustering coefficients (0.5-0.7). Modularity, as calculated by Louvain, was lower for the two continental networks (∼0.4 for both), and slightly higher for lake networks (ranging between 0.6 and 0.8) (Table 1). Among lake networks, the African crater L. Barombi Mbo was the most connected, showing the highest mean node degree (10.36) and graph density (0.09) and the lowest number of modules (8) (Table 1 and Fig. 1A). Of the two continental networks, the American network showed a slightly higher density (0.10) and mean node degree (14.84) compared to the African one (0.08 and 11.83).

**Table 1:** Global network topologies

|  | Dataset | Samples | N. of OTUs* | Nodes | Edges (+/-) | Mean Node Degree | Clust_coeff | Path_length | Density | N. of Modules | Modularity |
| --- | --- | --- | --- | --- | --- | --- | --- | --- | --- | --- | --- |
| **Continent** | America | 161 | 716 | 150 | 1113 (1083/30) | 14.84 | 0.69 | 3.34 | 0.10 | 17 | 0.44 |
|  | Africa | 116 | 637 | 149 | 881 (856/25) | 11.83 | 0.50 | 2.44 | 0.08 | 10 | 0.43 |
| **Lake** | Nicaragua | 24 | 504 | 147 | 483 (448/35) | 6.57 | 0.71 | 6.03 | 0.05 | 12 | 0.70 |
|  | Apoyo | 24 | 510 | 175 | 324 (321/3) | 3.70 | 0.61 | 2.68 | 0.02 | 32 | 0.82 |
|  | Apoyeque | 25 | 356 | 155 | 405 (370/35) | 5.23 | 0.60 | 5.05 | 0.03 | 18 | 0.57 |
|  | Xiloá | 43 | 494 | 158 | 364 (352/12) | 4.61 | 0.58 | 3.65 | 0.03 | 29 | 0.70 |
|  | Tanganyika | 73 | 597 | 162 | 276 (276/0) | 3.41 | 0.49 | 5.77 | 0.02 | 29 | 0.78 |
|  | Barombi Mbo | 43 | 431 | 122 | 632 (500/132) | 10.36 | 0.56 | 3.12 | 0.09 | 8 | 0.60 |

\* before filtering in CoNet

In terms of proportion of nodes assigned to a bacterial taxon, all lake networks showed a quite comparable profile, encompassing the same four major phyla (Proteobacteria, Firmicutes, Planctomycetes and Fusobacteria) (Fig. 1C), and six major families (Clostridiaceae, Pirellulaceae, Rhodobacteraceae, Fusobacteriaceae, Peptostreptococcaceae and and Lachnospiraceae), with comparable node frequency (Fig. S1 and S2). Proteobacteria contributed with the largest number of nodes, typically followed by Firmicutes. All lake networks, except for L. Tanganyika, showed a densely connected cluster of Firmicutes nodes (Fig. 1A, blue circles) which primarily belonged to two distinct families, Clostridiaceae and Peptostreptococcaceae (Fig. S1). Major differences in network composition were seen for L. Tanganyika, with its taxonomic composition dominating the pattern seen in the African network (Fig. 1C). This was characterized by a substantially lower proportion of nodes belonging to the phylum Bacteroidetes and Firmicutes, particularly of families Clostridiaceae (5% of the nodes) and Lachnospiraceae (no nodes) (Fig. S2), and a higher representation of Actinobacteria and Verrucomicrobia. The network taxonomic profile (proportion of node per phylum) only partly reflected the microbiota taxonomic profile (proportion of OTUs per phylum in the original input matrices, i.e. 774 OTUs, including only most abundant OTUs); the latter was characterized by a remarkably homogeneous pattern of phyla representation across lakes and continents (Fig. S3). This indicates that the network composition does not simply mirror the OTU diversity of the input dataset.

We next explored similarities among individual lake networks based on shared pairwise associations as measured by the Jaccard index (where edges are taken as observations) (Fig. 2A). A total of 1797 unique associations were obtained across all lake networks. The four American lakes resembled more each other than any of the two African lakes, while these, Barombi Mbo and Tanganyika, did not cluster together. Barombi Mbo network was more similar to any of the American lakes (6.9 to 11.2% of shared associations) than to L. Tanganyika (only 2.3% of shared associations), confirming findings based on node taxonomy (Fig. 1). The large majority of the two African lake associations were unique to each lake (i.e. 76% for crater lake Barombi Mbo and 87% for L. Tanganyika). Within America, the closest resemblance was observed between crater L. Apoyo and Xiloá networks (sharing 23.3% of their total unique associations), followed by similarity between Nicaragua and Apoyeque (21.6%). The four American networks shared overall between 44 and 55% of their associations with any other lake, with a total of 60 associations being shared across all lakes (Fig. 2B).

**Figure 2:**
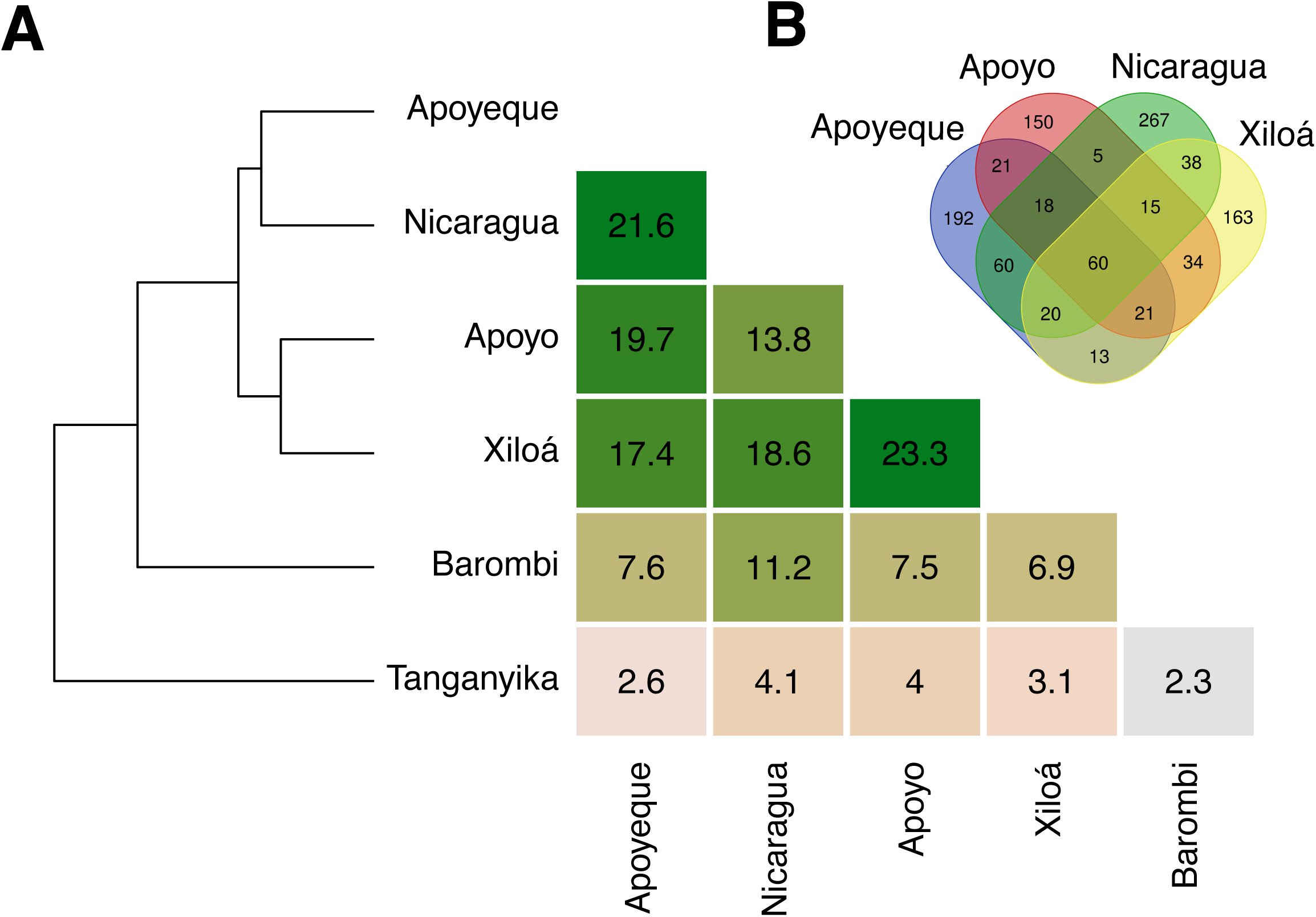
Distances among lake networks estimated by a Jaccard similarity index. (A) Heatmap of distance matrix values as proportion of shared OTU pairwise associations (i.e. edges) among lake networks and corresponding dendrogram. (B) The Venn diagram shows number of shared and unique associations among the four American lake graphs.

### Core microbial associations

To explore whether specific pairwise associations were conserved across the range of cichlid geographical distribution, individual lake networks were intersected to retrieve the shared component (Fig. 3). To this goal, we generated a core network for America, by intersecting the four lake-specific networks (Fig. 3A), and a core network for Africa, by intersecting lakes Tanganyika and Barombi Mbo networks (Fig. 3B). We finally retrieved associations common to all networks (Fig. 3C).

**Figure 3:**
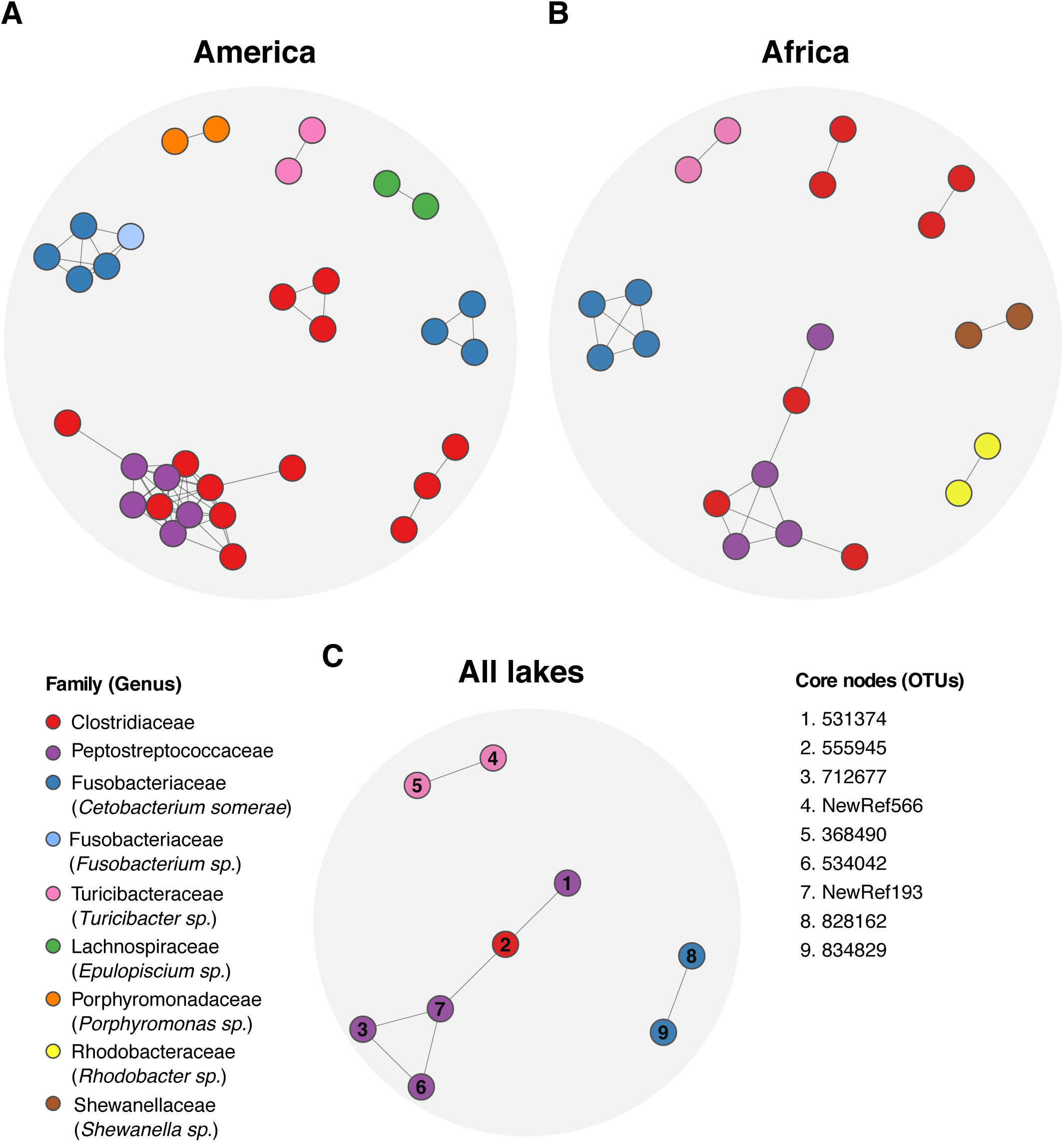
Core networks of common pairwise associations across geographical graphs. (A-C) Shared associations across the four American lakes (Nicaragua, Apoyo, Apoyeque and Xiloá; 60 edges, 32 nodes) (A), the two African lakes (Tanganyika and Barombi Mbo; 20 edges, 21 nodes) (B), and all lakes (7 edges, 9 nodes) (C). Nodes are colored according to the maximum taxonomic resolution achieved. Nodes in (C) are labelled by their OTU ID (right panel).

The four American lakes shared 60 pairwise associations involving 32 nodes/OTUs (Fig. 3A). These pairwises were structured into eight units (individual groups of nodes connected by at least one edge) comprising three phyla (Firmicutes, Fusobacteria and Proteobacteria) and six families. The major unit was composed of 12 nodes and included the two Firmicutes families, Clostridiaceae and Peptostreptococcaceae. All other units included associations among members of the same family (Clostridiaceae or Fusobacteriaceae), or the same genus (*Cetobacterium, Turicibacter, Epulopiscium* and *Porphyromonas*), except for one unit comprising two distinct genera, *Cetobacterium* (four nodes) and *Fusobacterium* (one node) from the family Fusobacteriaceae (Fig. 3A).

The African core (Fig. 3B) included 20 associations involving 21 nodes, structured into seven units. These encompassed the same three phyla characterizing the American core and six families (Fig. 3A). As for the American core, the major unit (seven nodes) included the two Firmicutes families, Clostridiaceae and Peptostreptococcaceae, while the other units involved associations among members of the same family (Clostridiaceae) or genus (*Cetobacterium, Turicibacter, Shewanella*, and *Rhodobacter*).

Overall, both American and African cores displayed a cluster of Firmicutes belonging to the two families Peptostreptococcaceae and Clostridiceae, a small cluster of *Cetobacterium somerae* OTUs and an association between a pair of *Turicibacter* OTUs (see also individual lake networks in Fig. 1A). The two continental cores differed for presence of *Porphyromonas, Epulopiscium* and *Fusobacterium* in the American core, and *Shewanella* and *Rhodobacter* for the African core (Fig. 3A and B).

The global network, obtained through the intersection of all lake-specific networks, encompassed seven pairwise associations among nine OTUs, structured into three units (Fig. 3C). These consistent associations can be considered as continent and lake-independent. The nine OTUs belonged to two phyla (Firmicutes and Fusobacteria), and four families (Peptostreptococcaceae, Clostridiaceae, Turicibacteraceae and Fusobacteriaceae). The first and largest unit comprised members of the families Clostridiaceae and Peptostreptococcaceae, the second unit included two members of the genus *Turicibacter*, while the third unit involved two members of the species *C. somerae*. Three of the nodes (OTU-555945, OTU-712677 and OTU-828162) represented core OTUs (found in 90% of the specimens according to (Baldo et al., 2019).

We note that the network intersections generated do not take into account indirect associations among OTUs, i.e. common nodes that are indirectly connected by means of few intermediate nodes (no direct edge connects the two nodes). Considering this caveat, the size of the core could be potentially larger.

### Diet-specific network features

We next focused our analysis on the L. Tanganyika dataset to explore key changes in microbial association patterns as a function of the cichlid diet. This lake assemblage encompasses the largest diversity of cichlid trophic diversity (our dataset including carnivores, scale-eaters, omnivores, planktivores and herbivores) and provides a reasonable number of samples per dietary category for data partitioning and reliable network inference. We tackled this goal through a double approach: first, we built separate networks for the two most representative diets, herbivores (H) and carnivores (C; includes scale eaters) and compared properties; in the second approach, we mapped diet-associated features (i.e. OTU abundances and indicator values) on the whole L. Tanganyika network.

The Tanganyika C and H networks showed remarkable differences in topology and node taxonomic representation (Fig. 4 for phylum and Fig. S4 for family levels), despite the use of a comparable sample size (*n* = 24 for C and *n* = 25 for H), similar OTUs richness in the input dataset (total number of observed OTUs = 460 for C and 542 for H) and same settings for network building (see Methods). The H network (156 nodes and 339 edges) was ∼8 fold larger than the C network (21 nodes and 70 edges) and taxonomically more diverse (Fig. 4A). Herbivores nodes belonged to nine phyla, the most conspicuous being the Proteobacteria (43%), followed by Planctomycetes (22%), Actinobacteria (15%) and Verrucomicrobia (11%); other phyla were poorly represented, including Acidobacteria and Chloroflexi (one node), Fusobacteria and Bacteroidetes (only three nodes each phylum), and Firmicutes (seven nodes) (Fig. 4A). Most represented families were Verrucomicrobiaceae, Rhodobacteraceae and Pirellulaceae (Fig. S4). Major hub nodes (according to normalized *betweenness*) involved members of the Proteobacteria (mostly of unknown families), Actinobacteria and Verrucomicrobia. In the C network (Fig. 4B), nodes encompassed a reduced diversity, represented by the same three phyla constituting the African core network: Firmicutes, Fusobacteria and Proteobacteria (see Fig. 3B). Most relevant hub nodes belonged to the Firmicutes families Clostridiaceae and Peptostreptococcaceae (Fig. S4).

**Figure 4:**
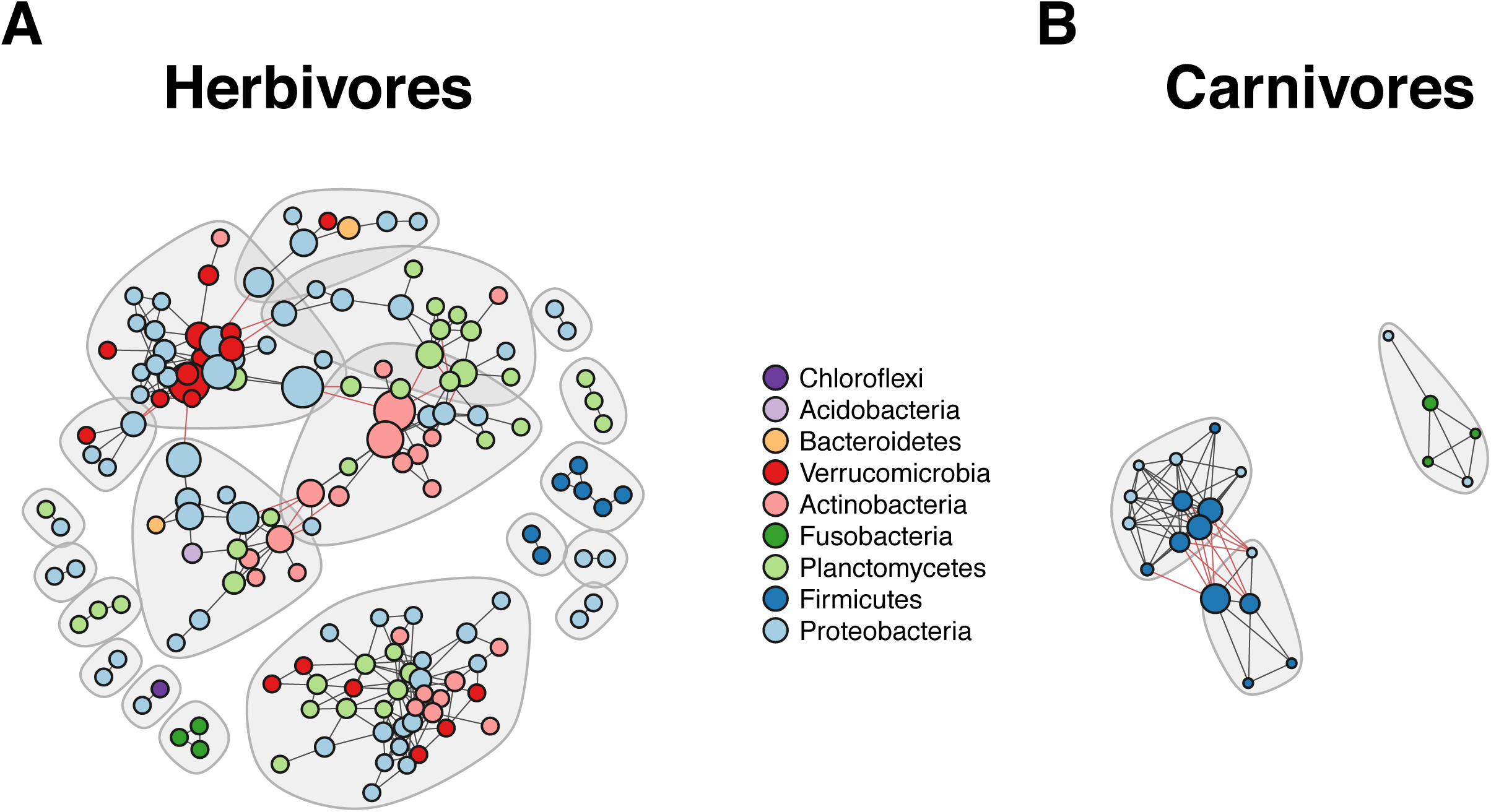
Diet-specific networks of L. Tanganyika herbivores and carnivores. Nodes are colored according to phylum and sized by betweenness values normalized by lake. Grey shades represent distinct modules (connected by red edges). Notice the higher complexity of the herbivore network (modules number, size and taxonomic composition) compared to the “minimal” carnivores network.

Modularity (according to Louvain algorithm) was ∼2.5-fold higher in the H (0.72) than in the C network (0.29). Herbivores microbial associations were structured into 19 modules, five of them encompassing more than 10 nodes each (Fig. 4, grey-shaded areas). In general, H modules encompassed members of distinct phyla and families, with interactions within modules typically involving members of the four most abundant phyla, Proteobacteria, Planctomycetes, Actinobacteria and Verrucomicrobia. The C network was structured into three modules only, each of them encompassing two phyla (Proteobacteria-Firmicutes and Proteobacteria-Fusobacteria). No bacterial associations were retrieved from the intersection of C and H networks, which shared only three isolated OTUs.

To characterize putative functional differences between the two diet-based networks we used FAPROTAX (see Methods). For both nets, the large majority of OTUs (between 50% and 70%) could not be assigned to any functional group, due to the low taxonomic resolution achieved, which rarely reach genus level despite the use of three distinct databases (RDP, SILVA and Greengenes) (results are not reported).

A diet effect on co-occurrence patterns was even clearer when we used a second approach, that is mapping diet-associated microbial features onto the L. Tanganyika network (Fig. 5). Both Wisconsin-transformed abundances and indicator values calculated for each of the four represented diets (carnivores, omnivores, planktivores and herbivores) were visibly displaced into different areas of the network (according to the optimal layout chosen, see Methods) (Fig. 5). There was a clear transition in the centroids of both estimates across the four diets within the network, with carnivores and herbivores representing the most discriminated, and omnivores and planktivores being closer to herbivores. This transition was also fundamentally supported by fish specimen mapping onto the network, with only few outliers (Fig. S5). Although the number of individuals per species do not allow for sweeping conclusions, the inspection of Fig. S5 suggests that there are differences among species and, intriguingly, differences tend to imply clusters of connected OTUs rather than random nodes. Diet-associated nodes in carnivores belonged to the same three phyla characterizing the carnivore network (i.e. Firmicutes, Fusobacteria and Proteobacteria, Fig. 4); these nodes tended to form few modules of small size and homogeneous taxonomic content (i.e. most associations occurring between members of the same phylum) and occupy a distinct network area with respect to other diets (Fig. 5). Both omnivores, planktivores and herbivores showed a relevant increase in number of diet-associated nodes and taxonomic diversity, with maximum diversity reached in herbivores. More importantly, most of these diet-associated nodes were not scattered throughout the network, rather they grouped into few large modules of heterogeneous taxonomic composition, mostly involving members of the phyla Proteobacteria, Verrucomicrobia, Planctomycetes and Actinobacteria (Fig. 5).

**Figure 5:**
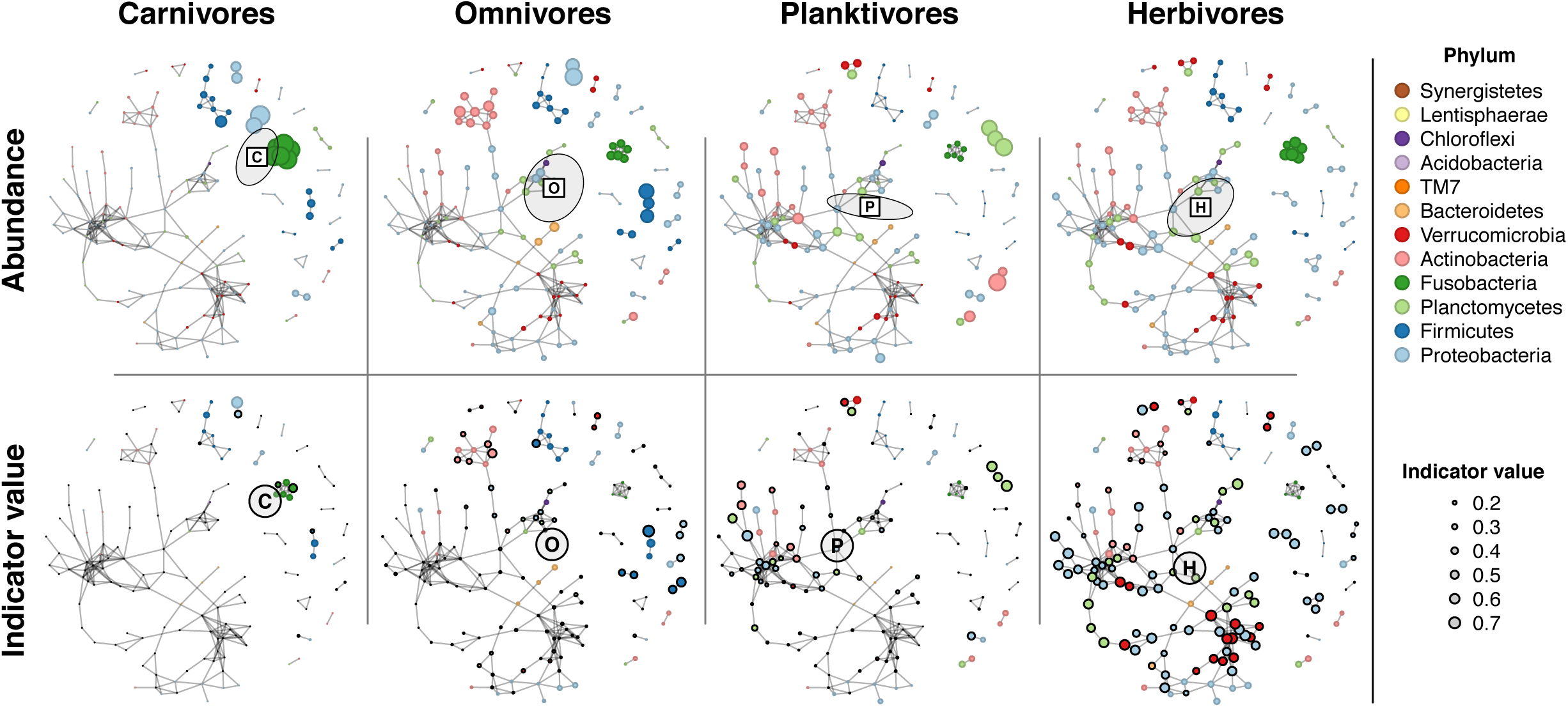
Diet contribution on the L. Tanganyika co-occurrence network. Four major diets (carnivores, omnivores, planktivores and herbivores) were mapped according to OTU abundance (after Wisconsin double transformation) (A-C) and indicator values (D-F). (A-C) For each diet, ellipses represent one standard deviation for the bivariate distribution of individual fish specimens, which are themselves positioned at their centroids (not shown for clarity, but see Fig. S5), calculated as average OTU coordinates weighted by transformed abundances. (D-F) Diet labels indicate centroids, weighted by significant indicator values. Circle nodes are sized by indicator value and framed in black color is significant at p < 0.05. Note that, whereas OTU abundances apply to individual specimens, OTU indicator values apply to diet groups. C: carnivores, O: omnivores, P: planktivores, H: herbivores.

## Discussion

We used the consortium of cichlid fishes, with their remarkable diversification, to explore gut microbial interactions through co-occurrence networks. Our main goals were to understand how much of the observed gut microbial diversity translate into robust microbe-microbe associations, whether there are conserved microbial co-occurrence patterns in cichlids, despite the scale of the host diversity (different species, ecology and geographical locations), and how community structures adjust following the host dietary shifts. To these aims we partitioned the dataset according to geographical and dietary variables, built individual networks and performed extensive comparative analyses.

### Cross-lake network comparative analysis reveals community conservation

Major findings of our study indicated that at least 21% (up to 43%) of total OTUs in individual cichlid lake assemblages (excluding the rare biosphere, see filtering in Methods) formed robust pairwise associations (based on concordance of four methods). These associations resolved into lake and continental scale-free networks that were largely comparable in terms of both topology (Table 1) and taxonomic content (Fig. 1). Major differences across geographical-based networks were primarily associated to L. Tanganyika, which hosts the most ecologically diverse of all cichlid assemblages (but see later for Tanganyika-specific aspects). The gut microbiota of all remaining lakes, including four American lakes (all crater lakes, except for lake Nicaragua) and the African crater lake Barombi Mbo, were all assembling into similar association patterns. The use of the same experimental approach and library protocol for all datasets, and sequencing in three distinct Miseq runs minimized technical artifacts in driving community structure similarity.

Persistent co-occurrence associations have been recently observed for free-living microbial communities, where a proportion of OTUs engage in similar interactions and form comparable communities in natural soils along large environmental gradients (Mandakovic et al., 2018). In host-associated microbiota, only a few sparse studies are beginning to explore the resilience and recurrence of similar association patterns across host populations (Jackson et al., 2018), species (Hegde et al., 2018), free *versus* laboratory-reared cultures (Hegde et al., 2018), and experimental treatments (Li et al., 2019). In humans in particular, specific interactions and network features (including modules) were found to be conserved across populations, suggesting that despite considerable intraspecific microbial variation and geographical distance, the microbiome tended to organize into stable interactions (Jackson et al., 2018). In our study we observed network taxonomic and topological properties conservation at a macroscale level, i.e. across gut communities encompassing different host species within a single family (the Cichlidae), and along a broad phylogeographic and ecological gradient. Specifically, gut community networks in all cichlid assemblages under study involved the same four major phyla and six families, with comparable representation across most lake networks (major differences were found for L. Tanganyika, Fig. 1C). American lakes formed particularly similar co-occurrence patterns: not only were their networks highly comparable in global topology (size, graph density, and clustering coefficient, Table 1) and node taxonomic content, but they also showed a considerable overlap in terms of specific OTU associations, sharing between 14 and 23% of all unique microbial pairs (edges) in any lake pairwise (Fig. 2). In particular, the crater lakes Xiloá and Apoyo represented the closest pair, a finding that is concordant with those obtained according to compositional aspects only (Baldo et al., 2019) and that might be driven by these cichlid’s minor dietary differences (being all largely omnivores) and recent lake diversification (Kautt et al., 2018). This suggests an interesting phylogeographic effect in shaping bacterial associations.

The overall conservation of association patterns observed, despite lake-specific features, supports the presence of common constraints in gut community assembling in cichlids. These constraints might be partly driven by putative conserved aspects of the cichlid gut environment, including physiology and immune system, that can favor specific microbe retention and persistence in the gut. In line with these findings, recent studies in fishes have shown a predominant role of host selection in shaping gut community composition, which appears as largely independent from the environmental microbial exposure (Yan et al., 2012, 2016). Moreover, the recent application of the ecological theory to the study of host-associated communities has provided a valuable theoretical framework to dissect the mechanisms driving community diversity and maintenance (Costello et al., 2012)(Miller et al., 2018). In fishes, for instance, deterministic rather than stochastic processes appear to guide the early onset of the gut microbiota assembling (Yan et al., 2016)(Yan et al., 2012), a scenario that could explain the similarities in cichlid association patterns observed across distinct lake assemblages. Experimental studies are clearly needed to validate the biological relevance of the observed co-occurrence relationships and to understand forces and ecological processes taking place in the cichlid gut.

Intriguingly, a small set of interactions was robust to all data partitioning, being consistently found in all networks (i.e. core, Fig. 3C). These seven recurrent pairwise associations, which can be considered as largely independent from major geographical variables, arranged into three distinct units. The largest of these units involved five OTUs belonging to two Firmicutes families, Clostridiaceae and Peptostreptococcaceae; these families formed consistent co-occurring pairs in all lake and continental networks where they resolved into large modules of interactions (Fig. 1A-B), although typically involving a distinct set of OTUs (except for the five core members). Whether these modules represent ecologically or functionally equivalent units across largely distinct systems clearly needs further investigation. Similarly, a consistent association was also retrieved between two *C. somerae* OTUs (Fig. 3A and B); this species is a vitamin B12 supplier (Tsuchiya et al., 2008) and represents the most conspicuous member of the cichlid gut, where it showed a systematic presence in all specimens (Baldo et al., 2017, 2019). Members of this species formed one/two conserved modules in each lake network, suggesting putative niche overlapping and/or diversification *in situ*. Finally, a recurrent pairwise interaction was also observed between two OTUs belonging to the genus *Turicibacter*, a common inhabitant of animal guts.

The nature and functional relevance of all these persistent associations remain unclear. An important aspect to consider is that the phylogeographical scale of network comparison in this study may be more relevant to detect association patterns at higher levels of bacteria hierarchical organization (from genera to families/orders), driven by ecological similarity within taxonomic group (Barberán et al., 2012), rather than among OTUs. A puzzling question is how the same co-occurring pairs of OTUs are established in these highly diverse lake assemblages. Vertical or restricted transmission of bacteria over time can also generate patterns of co-occurrence, and this is an avenue that is worth considering. Interestingly, both Peptostreptococcaceae and *Turicibacter*, here members of the cichlid core network, were shown to be highly heritable in mice and humans (Goodrich et al., 2016). Undoubtedly, the biogeography of these widespread OTUs and their ecological role need to be fully investigated if we are to understand their co-occurrence patterns in cichlids. This will require inclusion of additional replicated microbial community data, from the host and the environment.

### Microbial communities shift along dietary changes

How does the gut community structure rearrange in response to changes in dietary habits? To address this question, we excluded the major geographic effect (which constrains patterns of bacteria distribution) and focused our analysis on L. Tanganyika, which encompassed major dietary niches, exploring network topology and associations that were diet specific.

While there is extensive literature supporting a role of diet in changing the compositional aspects of the gut microbiota, diet-associated shifts in community association patterns have been little investigated (for an example outside humans, see (Li et al., 2019). We had previously shown that the taxonomic content of the cichlid gut microbiota was strongly affected by their dietary niche, with the herbivore-type gut microbiota being characterized by a significantly higher taxonomic diversity compared to the carnivore-type one (Baldo et al., 2017). Here we showed that this diversity resolved into highly distinct networks of interactions in the two extreme dietary categories (Fig. 4). In particular, the complex herbivore network, which was larger in size, modularity and taxonomic diversity, contrasted dramatically with the “minimal” network detected for carnivores. A similarly important community shift was observed by mapping diet-specific values onto the nodes of the whole Tanganyika network (Fig. 5). We found a significant increase in number and taxonomic diversity of abundant and indicator nodes from carnivores to omnivores to planktivores to herbivores, with a clear displacement of diet-based centroids along the dietary gradient. Interestingly, the same phyla that were shown to be significantly overrepresented in the African herbivores (Baldo et al., 2017) (Verrucomicrobia, Proteobacteria, Actinobacteria and Planctomycetes) were here found to form major clusters of positive associations (Fig. 4 and 5). The functional profiles of these clusters remain unresolved, as most of the OTUs involved are novel and show poor taxonomic resolution beyond the family level. Moreover, whether the coexisting microbial pairs observed are linked by metabolic interdependencies rather than shared niche preference cannot be presently discriminated through network analyses alone (Röttjers & Faust, 2018). Ongoing lab cultures are now targeting key community changes between cichlid specimens raised under high and low-fiber content diets (data in preparation). This could prove useful to test the predictive role of the hub taxa and association patterns here identified in driving gut community changes, as well as to explore bacteria functional relevance.

Additionally, fish accessibility to bacterial taxa involved in co-occurrence patterns needs to be investigated. Recent studies suggest that diet-specific taxa could be simply sourced from common dietary inputs. The macroalgal microbiota, for instance, is known to be significantly enriched in algal polysaccharide-degrading bacteria in comparison to the water column (Martin et al., 2015). This could potentially explain patterns of microbial co-occurrence across the diverse algal consumers from L. Tanganyika, a scenario that requires further investigation.

Altogether our study illustrates a powerful analytical framework to begin exploring community aspects of the gut microbiota along the scale of the host diversity. We showed that co-occurrence association patterns and particularly comparative network analyses of the gut microbiota can identify major constraints in microbial community assembly and maintenance, as well as specific co-occurrence relationships and hub taxa potentially involved in ecologically relevant transitions (Williams et al., 2014)(Faust & Raes, 2012)(Costello et al., 2012). By intersecting results from predictive network interactions and experimental trials, future studies will be directed to explore the strength of the cichlid-gut microbiota association, predict the outcome of community alterations driven by diet and ultimately help understanding the role of gut microbiota in cichlid trophic adaptation.

## Materials and Methods

### Sampled data and network generation

Samples corresponded to our previously published dataset (Baldo et al., 2019) and included a total of 278 specimens (42 species) sampled across nine lakes from Africa and America: the two African lakes, Tanganyika (*n* = 73) and Barombi Mbo (Cameroon, *n* = 43) and the seven Central American lakes, Apoyo (*n* = 25), Apoyeque (*n* = 25), Xiloá (*n* = 43), Masaya (*n* = 14), Asososca León (*n* = 13), Nicaragua (*n* = 29), and Managua (*n* = 12), all found in Nicaragua. Except for the large lakes Tanganyika, Nicaragua and Managua, all others are small crater lakes. African specimens encompassed multiple dietary niches, including carnivores, scale-eaters, omnivores, planktivores and herbivores, while American specimens are largely omnivores (see (Baldo et al., 2019) for details on diet assignment). Gut microbiota data was obtained through three distinct 16S rRNA Miseq runs using the same protocol for DNA extractions, library preparation and sequencing.

The original unrarefied matrix of OTU counts obtained after extensive filtering (see(Baldo et al., 2019)), corresponding to 3639 OTUs (16 083 429 total counts), was further filtered to remove OTUs contributing <0.005% of total counts across all samples. This step reduced the dataset to 774 OTUs and 15 467 721 total counts. To account for the heterogeneity of the data, the OTU matrix was then split into several datasets according to geographical (continent and lake) and ecological variables (diet) setting an arbitrary minimum number of samples for network generation (*n* = 20 fish specimens).

Each dataset (abundance matrix) was loaded into the CoNet ensemble app (Faust & Raes, 2016) available in Cytoscape v. 3.7.2 (Faust & Raes, 2016) for network generation (where rows are OTUs and columns are fish specimens). Settings were the same for lake and continental datasets. Specifically, the data was filtered for OTU occurrence across samples according to the minimum value suggested by the program, keeping the sum of filtered rows (row minimum occurrence = 38 for Africa and 53 for America, 24 for Tanganyika, 14 for Barombi Mbo, 8 for Apoyo, 8 for Apoyeque, 14 for Xiloá, 9 for Nicaragua). This step minimizes sparsity issues and false correlations due to double zero problem when computing similarity or distance coefficients using species presence-absence or abundance data (Faust & Raes, 2016; Weiss et al., 2016). In all cases, data was normalized by column (col_norm; entries divided by column sum) to reduce compositionality issues associated to different sampling efforts. Four correlation methods were chosen: Spearman, Pearson, Bray-Curtis dissimilarity and Kullback-Leibler dissimilarity, with automatic threshold set to retain 1000 edges (top and bottom) supported by all four methods (score = mean). Edge significance was tested through 1000 permutations and bootstraps (method = “brown”), retaining edges with merged p-values < 0.05 after Benjamini-Hochenberg’s correction (q-value).

To explore impact of diet on network topology and properties, the Tanganyika dataset was further split into herbivores (n=25) and carnivores plus scale-eaters (here grouped into the same category) (n=24) (for diet classification see (Baldo et al., 2017, 2019)). CoNet settings for dietary analyses were chosen to be more conservative: minimum OTU occurrence was set to 60% of the samples for both herbivores and carnivores, considering copresence only. For carnivores, we retained all positive edges resulting from the intersection of the four methods (quantile=1), providing they were less than 1000, while for herbivores we retained the top 1000 edges.

### Network analysis

Each network was loaded into R for all subsequent analyses. Network properties, including simple and complex parameters of global net and node topologies were estimated with *igraph* in R (Csardi & Nepusz, 2006). Number of modules, defined as clusters of nodes that form coherent structural subsystems of interacting units, and modularity (M) were measured according to the Louvain algorithm (Blondel et al., 2008). Hub nodes (OTUs) were explored by measures of normalized betweenness centrality with function *betweenness* in the *igraph* package. Distances among lakes networks were estimated by a Jaccard similarity index, taking edges as observations and building a distance matrix of edge presence/absence among lakes. Distances were graphically represented through a dendogram using the function *hclust* in base R.

The cichlid core was defined as the fraction of edges that were conserved across all lake networks. Shared edges were obtained through the function *intersection* in the *igraph* package.

The Functional Annotation of Prokaryotic Taxa (FAPROTAX) database (Louca et al., 2017) was used to assign functional categories to nodes (OTUs) from the carnivore and herbivore networks, according to the taxonomic classification provided by three databases: Greengenes, RDP and SILVA.

To map fish diet onto the Lake Tanganyika network, we started by choosing a network layout that maximized the separation among diets based on OTU abundance data after double row and column standardization (the so-called Wisconsin standardization). For a given network layout, each fish specimen contributing to the network construction can be plotted at the centroid of the abundance of its OTU abundances (calculated as the average of OTU layout coordinates weighed by their transformed abundance). This produces a map of fish specimens onto the OTU network layout. To obtain an appropriate network layout for visualizing differences across diets, we chose the Fruchterman-Reingold layout and searched for the random seed (among 10000 trials) that maximized the ration of the sums of squared distances among diets vs. within diets (i.e., an F analogue). Not all OTUs that are abundant (even after double transformation) discriminate well among diets. Therefore, in order to best visualize differences among diets, we mapped OTU indicator values onto the selected network layout. The indicator value of an OTU with respect to a number of groups (i.e., diets) is calculated as the product of its relative frequency and its relative average abundance in those groups (Dufrêne & Legendre, 1997). In other words, an OTU has high indicator value for a group if it tends to have high relative abundance in that group but not in the others, and within the given group, it tends to be present and relatively abundant in all members of the group. Indicator values were calculated using function *indval* in the *labdsv* package (Roberts, 2019), and mapped as node size onto the same layout as above, marking only OTUs with indicator values having p-value < 0.05 based on the randomization test in function *indval*.

## Data Availability

The microbiota data used in this study are available at Bioproject PRJNA531389 (American dataset) and PRJNA341982 (African dataset).

## ACKNOWLEDGMENTS

We thank Grégoire Maniel and Mariona Pajares Murgó for helping in the R scripts.

## Funding

This study was funded by the Agencia Estatal de Investigación (AEI) and the Fondo Europeo de Desarrollo Regional (FEDER), CGL2017-82986-C2-2-P to LB.

## Conflict of interests

The authors declare that they have no competing interests.

## Supplementary files

**Figure S1:**
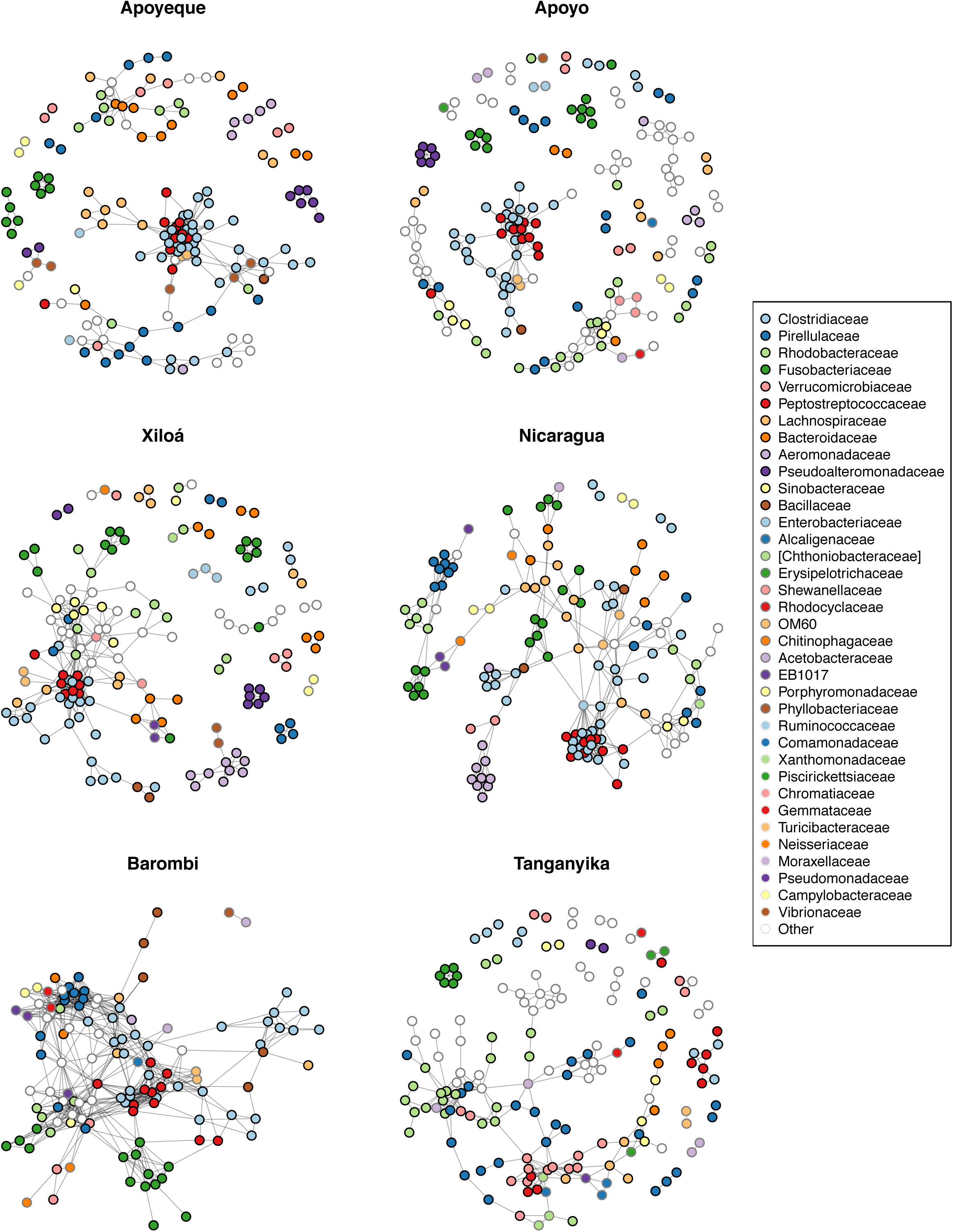
Individual lake networks with nodes colored by family level. For clarity, only the first 36 most abundant families are color-coded (remaining families are labelled as “Others” and shown in white). The same color is repeated across different families but in association to distinct border colors (i.e. black, gray and white).

**Figure S2:**
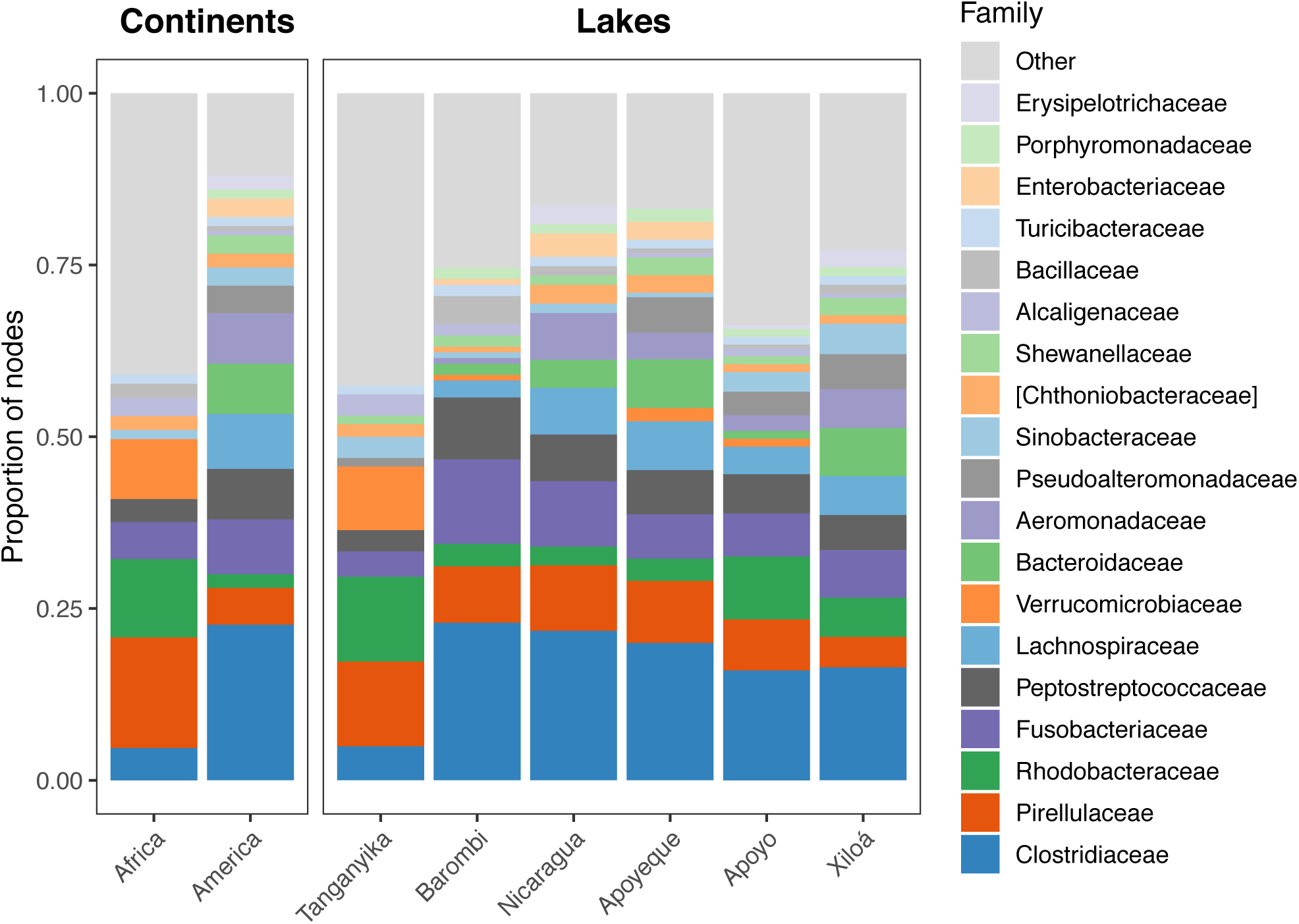
Taxonomic composition of continental and lake networks as proportion of nodes per family.

**Figure S3:**
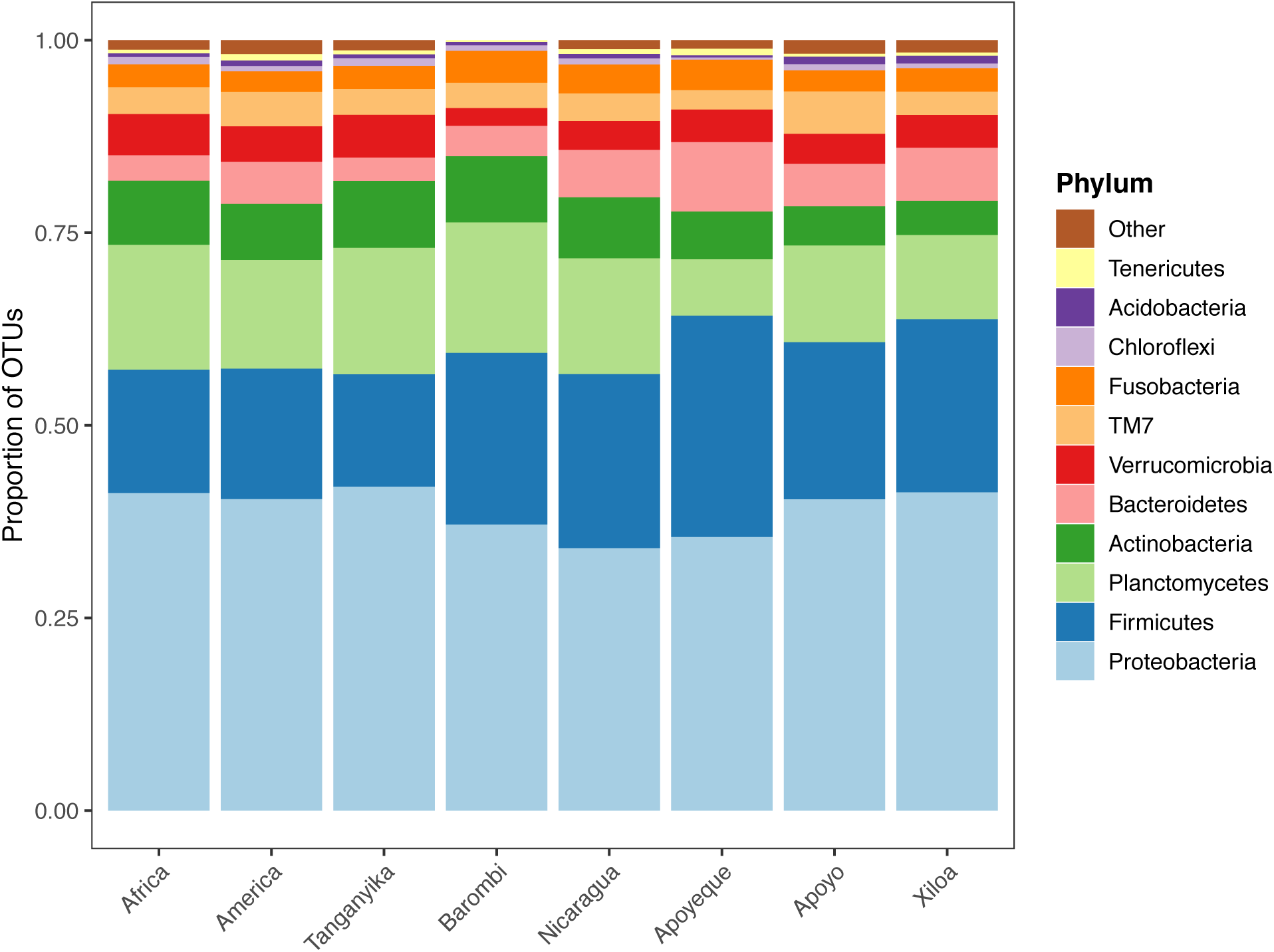
Microbiota taxonomic composition by lake and continental datasets expressed as proportion of OTUs present in the original input matrices (774 OTUs after filtering out low abundant OTUs).

**Figure S4:**
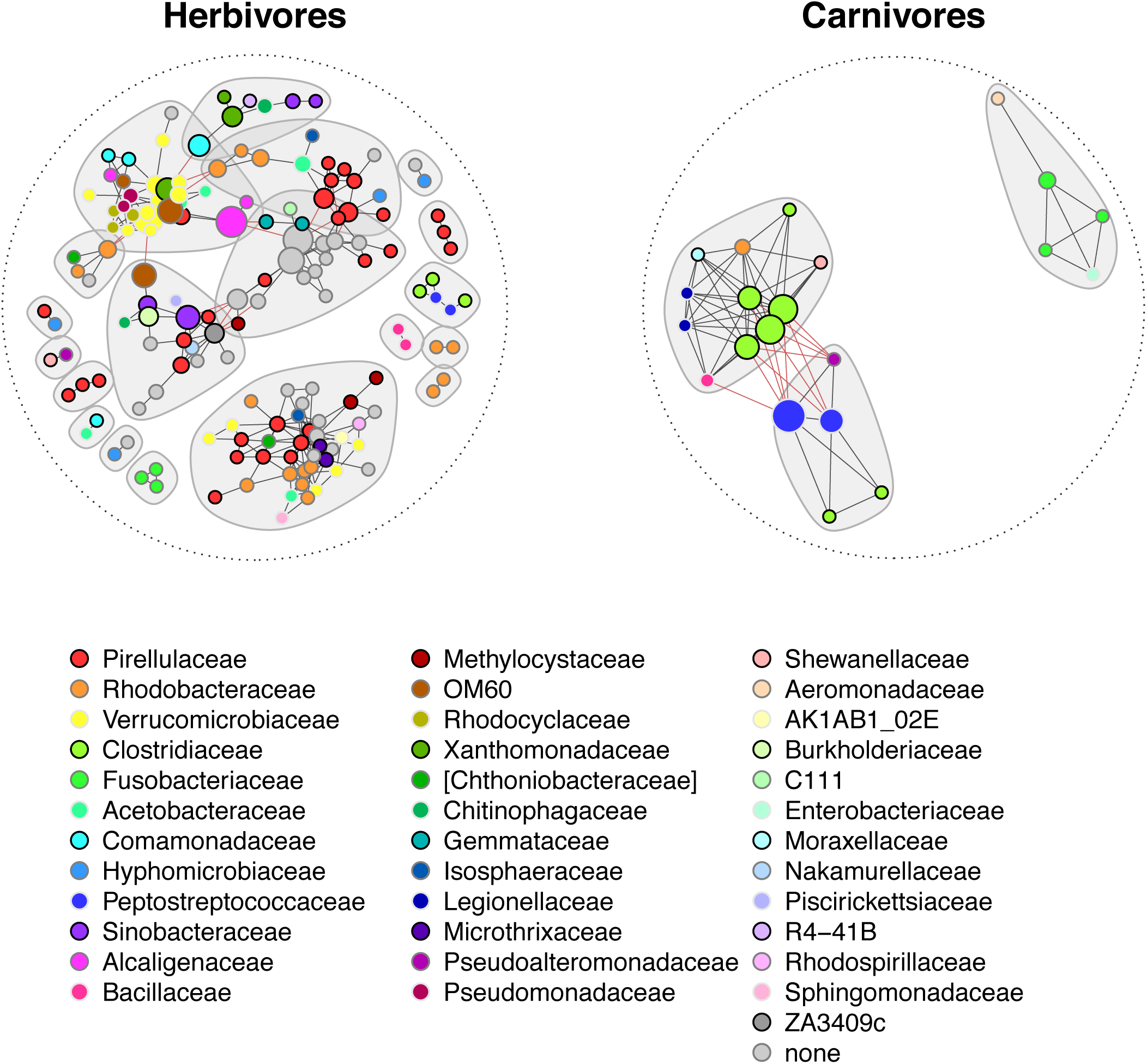
Diet-specific networks of L. Tanganyika herbivores and carnivores. Nodes are colored according to family and sized by betweenness values normalized by lake. Grey shades represent distinct modules (connected by red edges).

**Figure S5:**
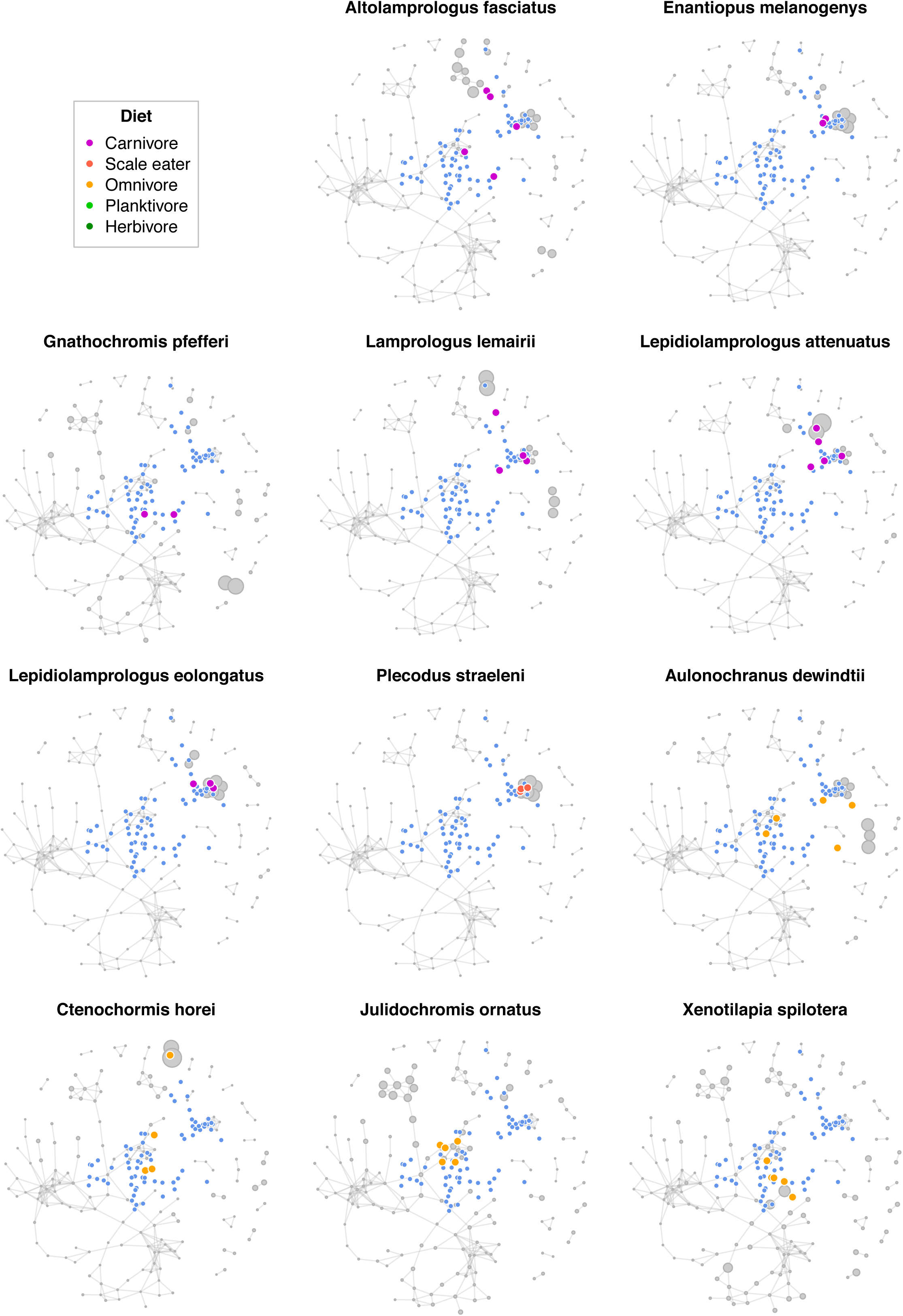
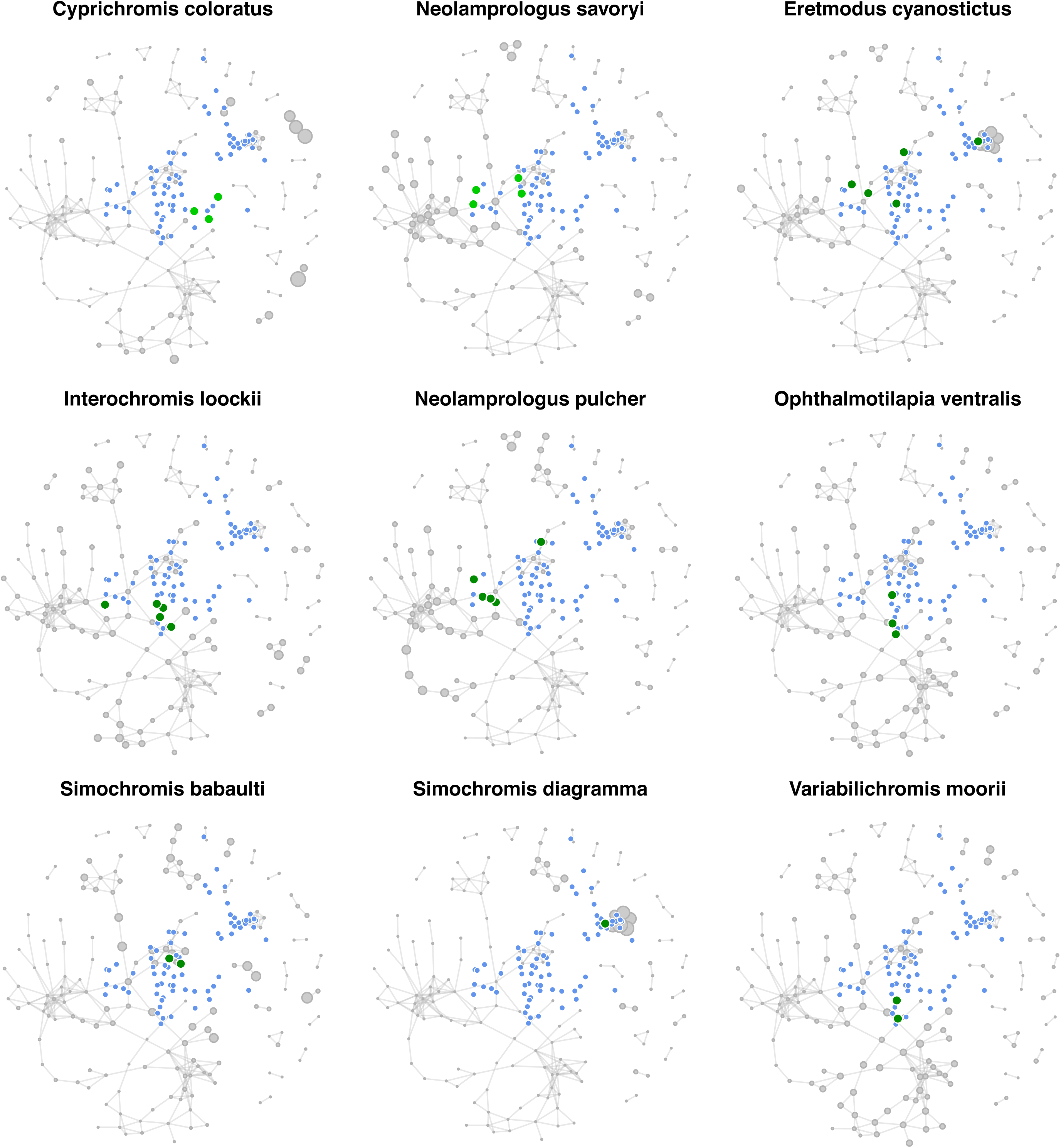
Mapping of individual cichlid species onto the Tanganyika network. The network layout is the same as in Fig. 5, with node sizes proportional to mean Wisconsin-transformed abundances in the specimens belonging to the given species. Colored points show the centroid position of each individual specimen within the network layout (calculated as mean OTU coordinates weighted by transformed abundances), and are color coded by diet. All remaining specimens are shown as blue dots for reference.

